# The Solvation of the *E. coli* CheY Phosphorylation Site Mapped by XFMS

**DOI:** 10.1101/2022.08.30.505875

**Authors:** Maham Hamid, Muhammad Farhan Khalid, Safee Ullah Chaudhary, Shahid Khan

**Affiliations:** Biomedical Informatics and Engineering Research Laboratory (BIRL), Lahore University of Management Sciences (LUMS), Lahore, Pakistan; SBA School of Science & Engineering, LUMS, Lahore, Pakistan; Molecular Biology Consortium, Lawrence Berkeley National Laboratory, Berkeley, USA

**Keywords:** solvent accessible surface area (SASA), mass spectroscopy, protein phosphorylation, protection factor (PF), MATLAB toolbox, structured water, protein fold energetics

## Abstract

The *Escherichia coli* CheY protein belongs to a large bacterial response regulator superfamily. X-ray hydroxy radical foot-printing with mass spectroscopy (XFMS) has shown that allosteric activation of CheY by its motor target triggers a concerted internalization of aromatic sidechains. We reanalyzed the XFMS data to compare polar versus non-polar CheY residue positions. The polar residues around and including the 57D phosphorylated site had an elevated hydroxy radical reactivity. Bioinformatic measures revealed that a water-mediated hydrogen bond network connected this ring of residues with the central 57D. These residues solvated 57D to energetically stabilize the apo-CheY fold. The abundance of these reactive residues was reduced upon activation. This result was supported by the bioinformatics and consistent with the previously reported activation-induced increase in core hydrophobicity. It further illustrated XFMS detection of structural waters. Direct contacts between the ring residues and the phosphorylation site would stabilize the aspartyl phosphate. In addition, we report that the ring residue, 18R, is a constant central node in the 57D solvation network and that 18R non-polar substitutions determine CheY diversity as assessed by its evolutionary trace in bacteria with well-studied chemotaxis. These results showcase the importance of structured water dynamics for phosphorylation-mediated signal transduction.

## 1. Introduction

The folding problem – the compaction of a 1D polymer to a 3D globule – is a fundamental problem in molecular biology. It has been known for some time that hydrophobic interactions form the protein core [1] - with globular proteins approximated as oil-drops coated by a polar shell that contacts the aqueous solvent. It is now appreciated that water-mediated interactions are also important for the protein fold [2], highlighted by the visualization of immobilized water molecules in high-resolution crystal structures. Studies have also revealed the role of structured water in protein allostery - the ability of proteins to change shape [3]. The role of structured waters is being extensively studied for drug design [4,5], with experiments, bioinformatic data mining and simulations to detect and characterize water dynamics in hydrogen bonded networks and internal cavities.

Protein folding and allosteric transitions are controlled by dynamic networks that integrate local transitions - the melting of α-helices, the rotation of aromatic sidechains and immobilization of surface loops [6] to trigger collective motions [7]. Time-resolved foot printing techniques are an important tool for mapping global protein transitions with single residue resolution based on the characterization by mass spectroscopy (MS). X-ray hydroxy radical foot printing (XF) with mass spectroscopy (XFMS) is one such method that monitors the solvent accessibility of residue sidechains to map conformational transitions in solution [8]. XFMS studies have shown that residues in proximity to structured water have anomalous oxidation rates that are elevated relative to residues in contact with bulk water (reviewed by [9]). These studies have targeted membrane G protein coupled receptors (GPCRs) and channel proteins where the anomalous reactivity is readily correlated with the presence of the few buried water molecules seen to be near the affected sidechains [10-12]. For soluble proteins, the temperature dependence and O^16^/O^18^ exchange has distinguished between bulk versus structured water residue oxidations. Oxidation of a cytochrome-C (Cyt-C) residue (67Y) hydrogen-bonded to water in the crystal had a markedly longer response time to fast O^16^/O^18^ water exchange than other Cyt-C residues [13]. Since this work, the interpretation of XFMS dose response curves has advanced by the development of protection factors (PFs) [14,15], a quantitative metric for the relation of oxidation rate to solvent accessibility [12].

We wondered whether the analysis of protection factors (*PF*s) might offer a rapid diagnostic to separate bulk and structured water oxidations in soluble proteins where the large number of protein-associated waters in the crystal structures makes a straightforward correlation between anomalous reactivity and water proximity difficult. We reworked XFMS data acquired previously on the *Escherichia coli* CheY bacterial response regulator [16] to test this idea. CheY, a 14 kilodalton protein with a α/β flavodoxin-type fold, is the founding member of a large aspartyl phosphate response regulator superfamily involved in diverse intracellular signal transduction circuits [17]. The CheY crystal structures have revealed an extensive hydrogen bonded protein-water network modulated by activation by the N-terminal peptide of the flagellar motor protein FliM (FliM_*N*_) [18-20].

SPECTRUM is an open-source software suite for high-throughput proteoform identification [21]. It’s extension, *SPECTRUM-XFMS*, a custom high-throughput algorithm, was used for the XFMS analysis. Recently developed bioinformatic algorithms, in addition to inspection of the crystal structures, were utilized to infer how the water associated with the polar residue positions detected by XFMS affected the energetics and the hydrogen-bonded network of the protein fold. We found that these residues formed an ion ring around the 57D phosphorylation site. The ring was reduced upon activation consistent with water depletion due to the stabilization of the hydrophobic core by concerted internalization of aromatic sidechains [16]. The evolutionary trace of CheY proteins specifically involved in chemotaxis revealed that non-polar substitutions in one residue position (*E. coli* 18R), a central node in the hydrogen-bonded network, were central to diversity of the CheY chemotaxis proteins. These results advance knowledge of CheY signal transduction and, more generally, develop XFMS methodology as a diagnostic tool for structured waters.

## 2. Results

### 1. Polar residues have lowered protection against hydroxy radical oxidation

The *E. coli* apo-CheY (3CHY.pdb) and CheY13DK106YW.FliM_N_ (1U8T.pdb) crystal structures, henceforth cited as Mg^2+^.free *CheY*_*I*_ and *CheY*_*A*_ respectively, reveal an extensive hydrogen bonded network. The CheY 57D phosphorylation site is ringed with polar residues (E, D, K, R, N) for coordination of the magnesium ion and the aspartyl phosphate. The XFMS oxidation dose response plots for these 10 residues, determined in Mg^2+^ buffer solutions for both states (**Materials & Methods**) are shown in **Figure 1**. The oxidation of 9 other ring residues was detected for one state alone. One residue (75D) external to the ring was oxidized in both states. Its oxidation rate increased in the active form as opposed to the rate decrease for the majority of the 57D ring residues. The *E. coli* protein sequence covered (89%) by the tryptic peptides submitted for MS analysis included 35 of the 42 total CheY (E, D, K, R, N) residues. The oxidation of the remaining 15 (E, D, K, R, N) residues was not detected.

**Figure 1:**
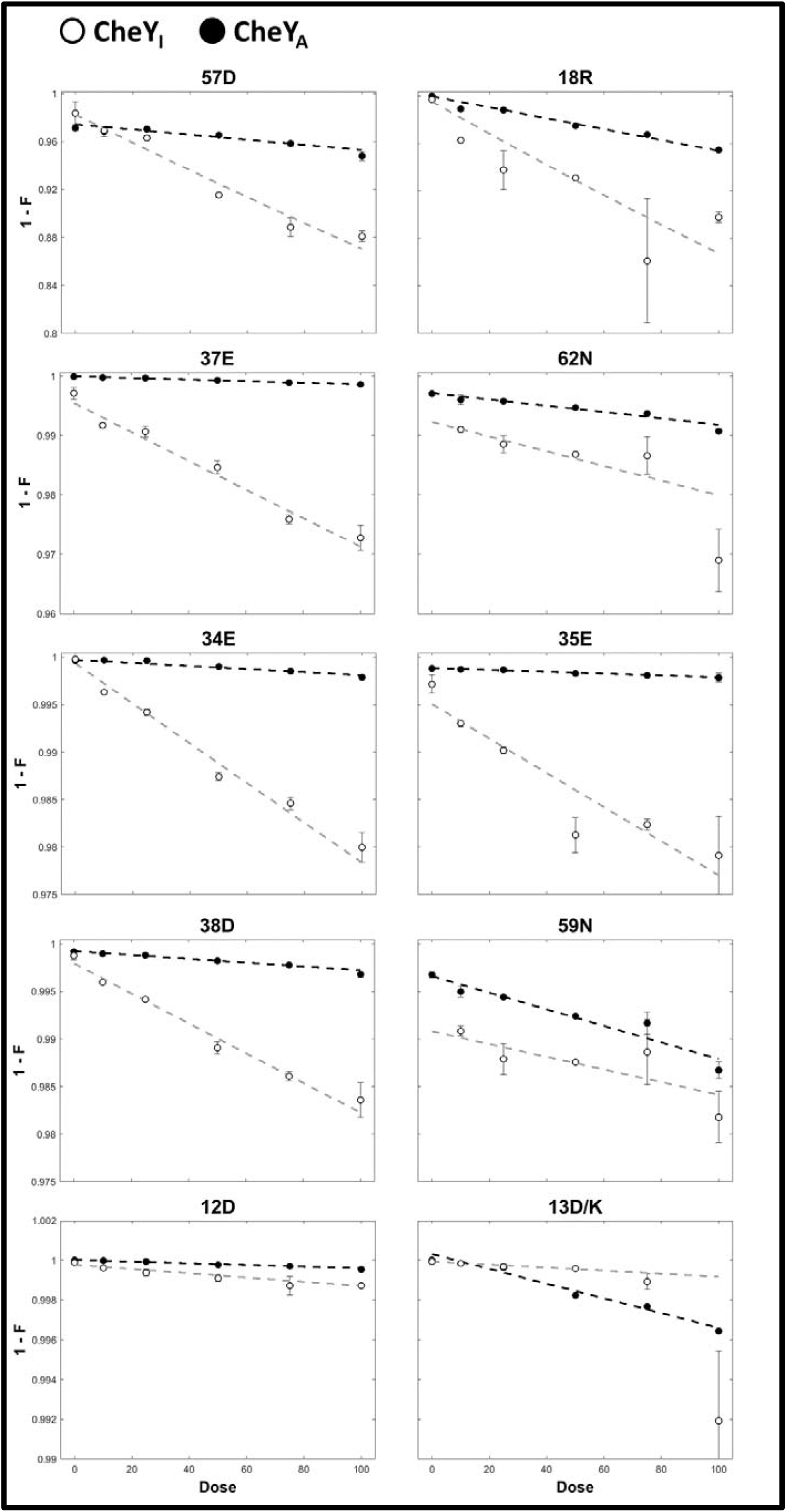
XFMS dose-response curves for residues around the 57D phosphorylation site. Oxidation rates were computed from the slopes of single exponential fits, weighted by reciprocal of the standard error, for all ten residues, including 57D. The CheY_A_ rates (black dashed lines) were lower those for CheY_I_ (grey dashed lines) in all cases, except for 13D/K and 59N. The intrinsic reactivity to hydroxy radical oxidation is greater for K (2.2) than D (0.42).

The solvent accessible surface area (SASA) - Log (*PF)* regression was constructed from the linear best-fit to *CheY*_*I*_ non-polar methionine and aromatic residue oxidations plus the *CheY*_*A*_ aromatic residue oxidations alone, as methionine was replaced by selenomethionine in the *CheY*_*A*_ construct used for crystallization (**Figure 2**). The Pearson’s correlation (-0.6) was comparable to that reported previously [16]. The corresponding values for the polar sidechains were then superimposed on the plot for comparison. They partitioned into two distinct sets demarcated by the lower 95% confidence limit to the best-fit relation for the non-polar sidechains. One set (12D, 13D/K, 37E, 59N, 62N, *CheY*_*I*_ (27E, 64D, 89E, 91K, 106K), *CheY*_*A*_ (38D, 118D, 119K) followed the (SASA) - Log (*PF)* relation obtained for the non-polar residues. The second set with Log (*PF)* values below the lower limit consisted of three residues 18R, 57D (phosphorylation site) and 75D. The *CheY*_*I*_ residues 34E, 35E, 38D, 41D, 94N also belonged to this set.

**Figure 2:**
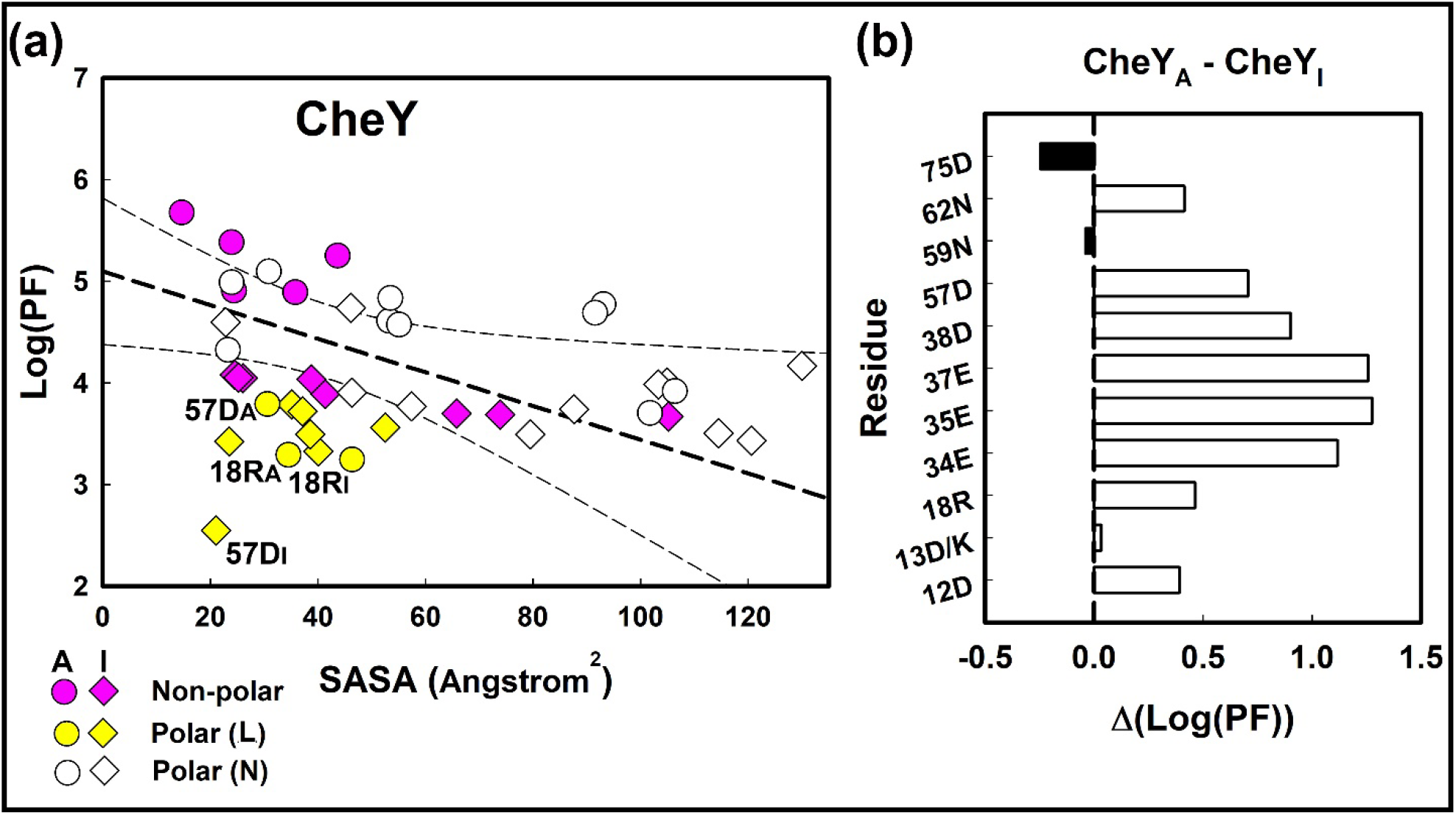
PF differences between residue types and activated states. **(a)**. The solvent accessibility of non-polar residues (magenta symbols) varies linearly with SASA. The least-square best fit linear regression (thick dashed line) and 95% confidence limits (thin dashed lines) are shown. Most polar residues (white symbols (N)) follow the best-fit relation, but a significant fraction (yellow symbols (L)) have anomalously high solvent accessibility (low Log (PF)) A = CheY_A_, I =CheY_I_. **(b). The activation dependent change in polar residue Log (PF) values**. There in a marked positive Log (PF) (Δ(CheY_A_ - CheY_I_)) increase for the 34E, 35E, 37E, 38D residue quadruplet ringing 57D, modest increases for 12D, 18R, 57D, 62N, and no significant change for 13D/K, 59N. The Δ(CheY_A_ - CheY_I_) Log (PF) for 75D outside the 57D solvation network has a negative value.

The non-polar *CheY*_*A*_ residues spanned a smaller SASA relative to their *CheY*_*I*_ states, in line with activation-dependent burial of aromatic sidechains. The non-polar sidechains had increased protection in *CheY*_*A*_ relative to *CheY*_*I*_ (**Figure 2b**). There was no change (ΔSASA = - 1.0±6.9) for the polar residues, except the change due to the 13D/K substitution, in contrast to the aromatic residues (ΔSASA = 28.3±10.8).

### 2. CheY activation stabilizes the 57D phosphorylation site

Next, we analyzed the PDB structure files to study the role of local waters in the deviation from the (SASA) - Log (*PF)* relation established for the non-polar residue positions.

First, the energetics of the contacts between the oxidized residue positions was estimated by Frustratometer2 based on knowledge of the local dielectric and packing density in the CheY fold (**Figure 3**). Energetically stressed contacts with *CheY*_*I*_ 18R, 34E, 38D, 37E, 41D, 94N (low PFs) ringed the 57D phosphorylation site. The oxidized residue positions (normal PFs) completed the ring with 75D the sole exception. The energetics of a Mg^2+^-bound *CheY*_*I*_ state (2CHE.pdb [22]) were also analyzed. The Mg^2+^ ion did not notably alter the ring of energetically stressed contacts. In contrast, the ring was reduced in *CheY*_*A*_. This reduction correlated with the reduction in the number of low PF residue positions to 18R consistent with water depletion from the more hydrophobic core due to burial of aromatic residue sidechains. The decrease in frustrated contacts provides an independent metric to the ionizable residue electrostatics used previously [16] to document the stabilization of the CheY core upon activation.

**Figure 3:**
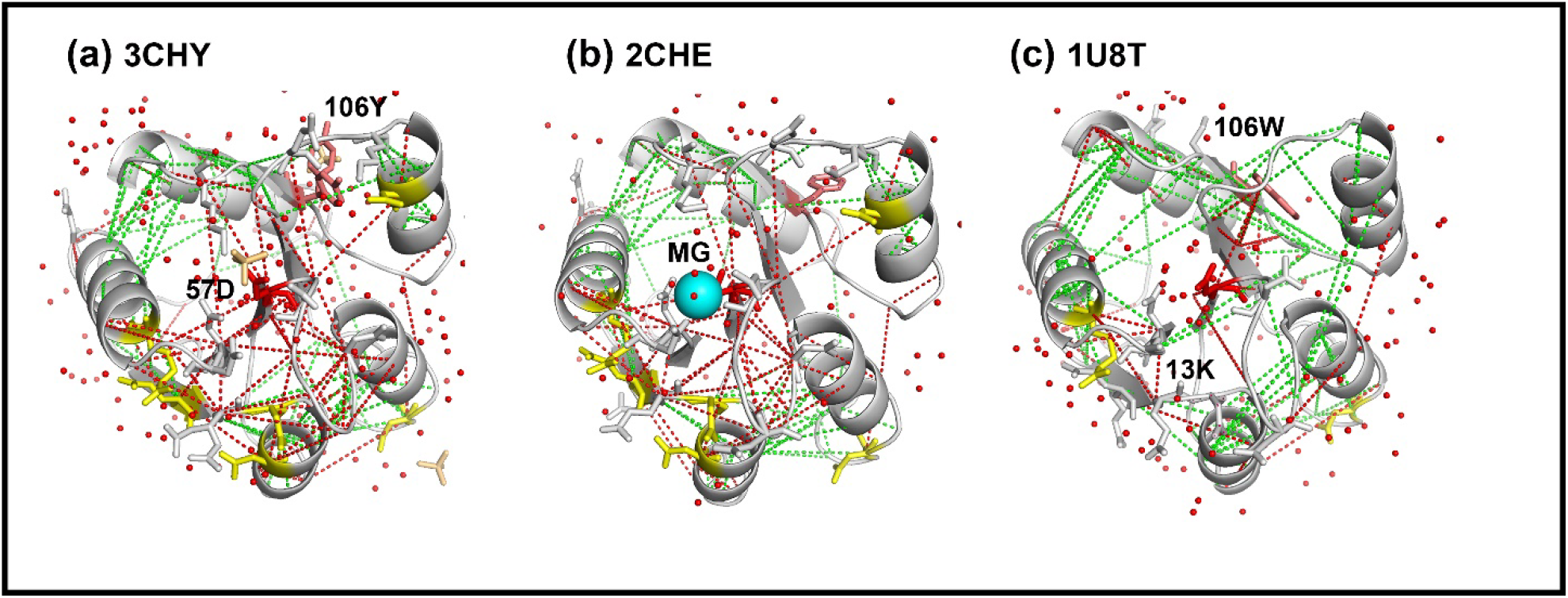
Activation dependent change in fold energetics. The Frustratometer2 algorithm computes residue contact energies relative to a decoy contact distribution with a coarse-grained AWSEM force-field. The PDB structures with superimposed high (red lines) and low (green lines) contact energies relative to the decoy distributions are shown. **(a) Apo-CheY**. High-energy contacts radiate out from 57D (red sidechain) to surrounding residues that include residue positions detected by XFMS (stick sidechains) with anomalous low (yellow) or comparable (white) Log (PF) values to non-polar residues (see text). 106Y sidechain (salmon). **(b) Mg**^**2+**^ **bound apo-CheY**. Magnesium ion (cyan sphere). **(c) Phospho-mimic 13DK**,**106YW CheY**. Activated by bound FliM_N_ (not shown). XFMS residue positions in (b), (c) as in (a). There is a reduction in the high energy contacts of 13D/K, 106YW CheY 57D with surrounding residue positions relative to the apo-CheY forms. Waters (red dots.).

Frustratometer2 does not report on water positions, but infers water mediated contacts based on the protein packing density. It assessed that all polar residues detected by XFMS had water-mediated contacts with their neighbors that included 57D in several cases. The energetically stressed contacts may have been maintained for formation of the aspartyl phosphate, with the waters acting to bridge negatively charged sidechains. In the phosphomimic *CheY*_*A*_, the 13D/K substitution would reduce the water requirement for such purpose.

### 3. Graphical analysis of the protein-water hydrogen bonded network

There was a qualitative correlation between the Frustratometer2 inferred water-mediated contacts with water positions seen in the crystal structures, but the coarse-grained AWSEM (associative memory, water mediated, structure and energy model) force-field on which the algorithm is based precluded a rigorous analysis. Therefore, we analyzed the network architecture of the crystal structures with Bridge2 to better characterize the local solvent density. The Bridge2 algorithm, developed since publication of these structures, is a graphical tool for representation and analysis of the full-atom protein-water hydrogen bond network. The bonds included intervening water molecules [19,20]. There are 147 and 136 water molecules associated with the CheY monomer in the Mg^2+^ free, 1.7 angstrom resolution 3CHY.pdb and 1U8T structures respectively. In addition, Mg^2+^ bound structures of a *CheY*_*I*_ form (2CHE.pdb (1.8 angstrom resolution, 99 waters [22])) and a beryllium fluoride (BeF_3_) *CheY*_*A*_ *F*_*3*_ form in complex with CheZ peptide (2FQA.pdb (2.0 angstrom resolution, 80 waters [23])) were analyzed. We obtained maps of their protein-water networks characterized by their centrality (**Figure 4**). The networks were superimposed on the 3D structures and displayed as a 2D map. Inspection of these representations yielded insights into the rearrangement of the hydrogen-bonds involved in allosteric activation of CheY.

**Figure 4:**
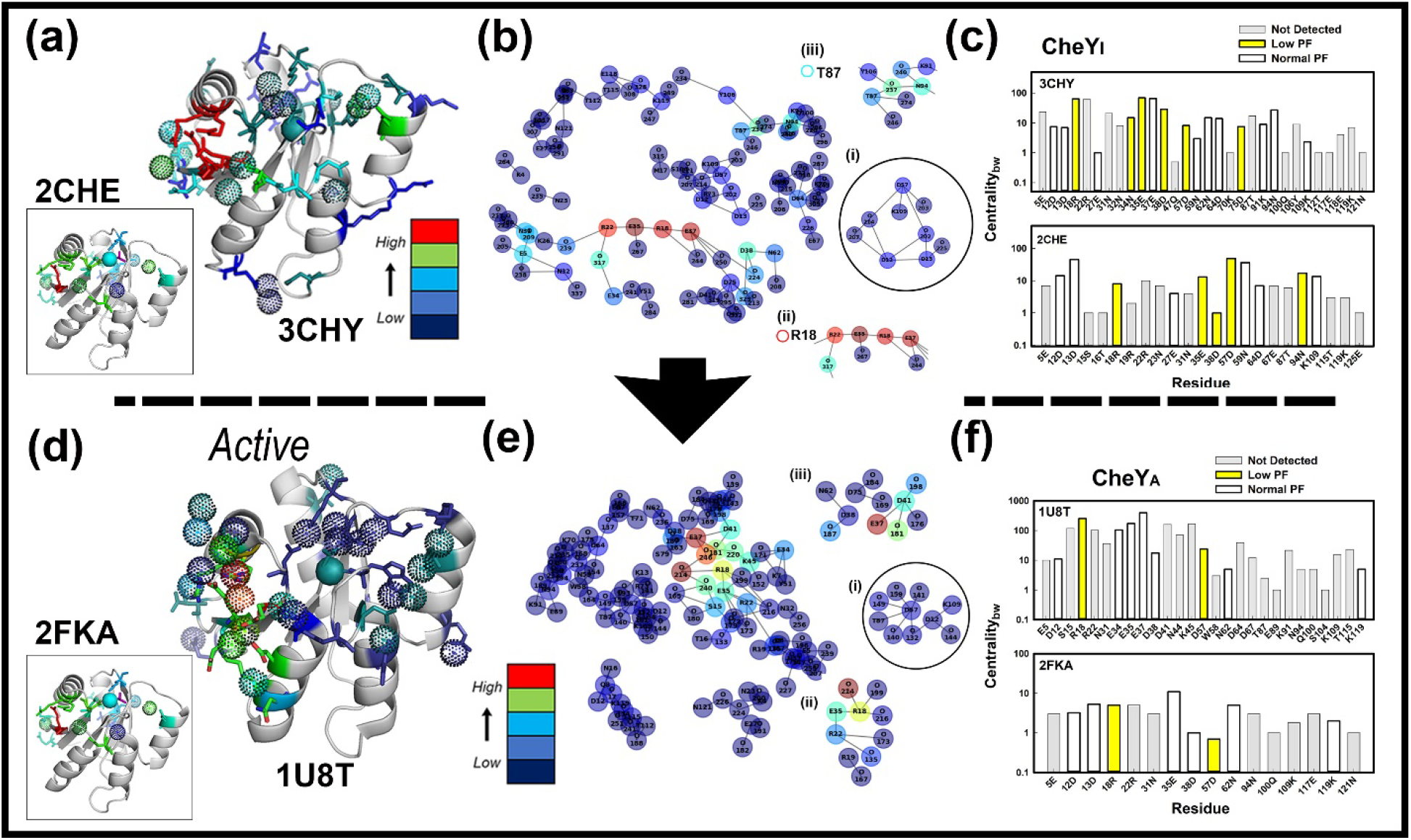
Changes in the Bridge2 CheY hydrogen-bonded protein-water network upon activation. Bars = Color coded C_bc_ values (Blue (Low) -> Red (High)). **(a) – (c). Inactive apo-CheY. (a) The C**_**bc**_ **nodes mapped onto the 3D structure**. Residues detected by XFMS (stick sidechains). Water (dotted spheres). Mg^2+^ (cyan spheres), BeF_3_ (magenta sticks). **(b) The 2D map of the network**. Local nodes **(i)** 109K -> 57D (circled [19]) **(ii)** 18R. **(iii)** 94N -> 106Y. (**c) The C**_**bc**_ **nodes targeted by XFMS detected residues C**_**bc**_ **nodes mapped onto the 3D structure**. Residues detected by XFMS and waters (dotted spheres) as in (low (yellow), normal (white)). **(d) – (f). Activated CheY13DK106YW in complex with FliM**_**N**_. **(d) The (a). (e) The 2D map of the network**. Local nodes **(i)** 109K -> 57D (circled [19]) **(ii)** 18R. **(iii)** 37E -> 41D, 38D -> 62N,75D **(f) The C**_**bc**_ **nodes targeted by XFMS detected residues** (anomalous (yellow), comparable (white)). Bridge2 residue nomenclature, where type precedes position, is different from nomenclature used elsewhere in the text.

Inspection of the 3D maps (**Figures 4a, d**) superimposed on the crystal structures showed both protein and water nodes clustered around, and including, 57D. A structurally conserved dome of water molecules above 57D was a feature of this water map. The prominent 18R residue node connected the 57D network, with nodes mapped on the α_1_-β_2_ surface.

The Bridge2 2D maps (**Figures 4b-c, e-f**) reported three features of interest. First, the algorithm faithfully documented the published [19], hydrogen-bonded rearrangements triggered by allosteric activation. Second, the dominant clusters in both cases were centered around 18R / 35E. Additional clusters were the water-mediated 87T-94N-106Y cluster in *CheY*_*I*_ 3CHY and the 37E, 41D, 38D-62N, 75D clusters in *CheY*_*A*_ 1U8T. Third, the anomalous, low (PFs) residue positions (ALPFs) revealed by the XFMS experiments targeted the centrality *(C*_*bc*_) profiles of the Bridge2 mediated protein-water networks. The *C*_*bw*_ values report high connectivity within the protein-water network not local solvent density, but we hoped that high values reporting hydrogen bonded clusters would be correlated with solvent density. The *CheY*_*I*_ 3CHY network (15 water, 30 (detected 17) amino acid) residue node *C*_*bc*_ values were 31.3±9.6 (low), 15.1±6.1(normal), 11.1±4.4 (not detected). The 2CHE network (7 water, 23 (11 detected) amino acids) residue node *C*_*bc*_ values were 17.5±8.2 (low), 19.9+6.8 (normal), 4.3±0.9 (not detected). The merged *CheY*_*I*_ networks had a (*Detected*/*Undetected*) *C*_*bc*_ value ratio of 2.6±0.6. The residue *C*_*bc*_ node values for the 1U8T network (35 water, 27 (detected 9) amino acids), were 137.9±114.1 (low), 101.6+54.9 (normal), 43.8±13.0 (not detected). The residue *C*_*bc*_ node values for the 2FKA network (5 water, 16 (detected 8) amino acids) were 2.8±2.1 (low), 4.6+1.4 (normal), 2.7±0.9 (not detected). The merged *CheY*_*A*_ networks had a (*Detected*/*Undetected*) *C*_*bc*_ value ratio of 2.4±1.2. The separation of the ALPFs 18R and 57D from other residues detected by XFMS for *CheY*_*A*_ may not have statistical significance. Nevertheless, the (*Detected*/*Undetected*) *C*_*bc*_ ratios for both *CheY*_*I*_ and *CheY*_*A*_ were significant, consistent with a correlation between *C*_*bc*_ and solvent density.

### 4. The ALPF 18R is important for chemotactic diversity

The 57D, 12D and 13D residue positions are important for phosphorylation, hence chemotaxis[17]. There are no known residue substitutions to our knowledge that affect chemotaxis in the ALPF 18R or the *CheY*_*I*_ ALPFs 34E, 35E, 37E and 38D. Therefore, we used EV trace to assess the importance of these residues to chemotactic function, in particular residue 18R prominent is mediating both connectivity and stabilization of the CheY core.

The EV-Trace for *E. coli* CheY matched the trace deposited in UniProt for the closely related *Salmonella typhimurium* CheY. Each trace collected 421 sequences for a multiple sequence alignment (MSA) and phylogenetic tree construction (**Figure 5a**). The rvET scores combined both positional conservations, as determined by the MSA, and evolution, as determined by the phylogenetics. The scores for the 12 top ranked residue positions (10% of the protein) are listed in **Figure 5b**. The residue 18R ranked tenth on this list. Five of the other remaining residues typically determine fold architecture and dynamics. The prolines (61P, 110P) disrupt α-helix formation and the glycines (39G, 65G, 110G) form flexible hinges. Mutational studies have established the importance of 109K and 87T for chemotaxis, in addition to 12D, 13D and 57D [24]. Thus, 18R together with 60M are the only residues in the list whose functional relevance is not known.

**Figure 5:**
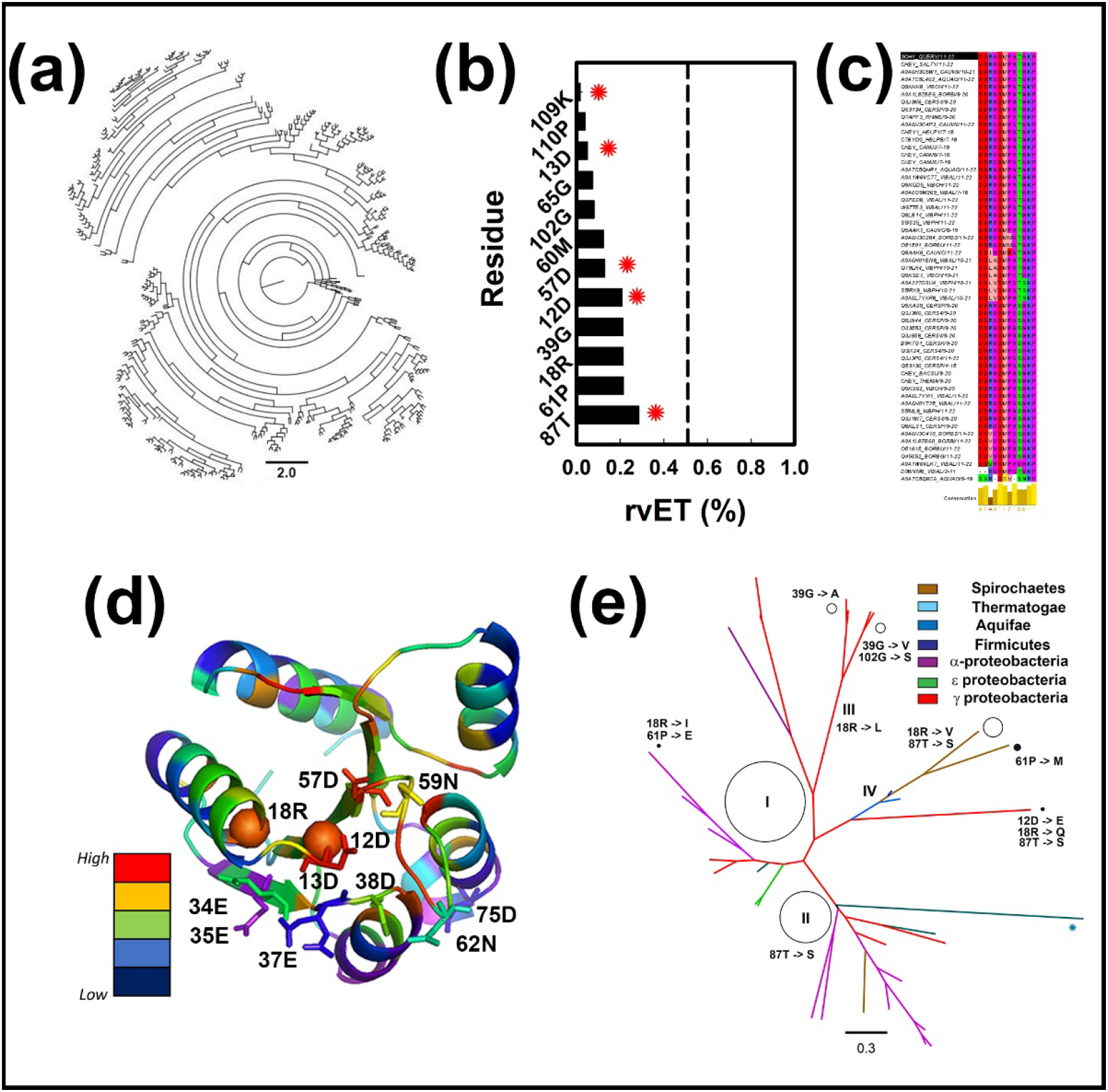
The evolutionary importance of ALPF 18R in CheY evolution. (a) The phylogenetic tree for E. coli CheY. Based on 443 sequences used by EV-Trace. Scale bar = 2.0. **(b) Top (10%) residue positions**. The E. coli CheY residue positions were ranked by rv.ET score. Asterisks denote residues known previously to be important for chemotaxis. **(c). Residue conservation of the top rv.ET ranked positions**. From an MSA of 56 sequences from 12 distinct species. **(d) The 3D CheY structure color-coded by ev.ET score**. The residues oxidized by XFMS are labelled (stick sidechains). Spheres mark the residues 12D and 18R (spheres) important for CheY evolution under selection for chemotaxis. **(e) The phylogenetic tree constructed from the MSA of CheY chemotaxis proteins**. 56 CheY sequences. The non-polar 18R substitutions localize at the internal tree nodes. Roman numerals denote major branches. Circle diameter = branch size. Note difference in scale with the EV-Trace tree (a).

We parsed out CheY sequences dedicated to chemotaxis from the large and functionally diverse bacterial response regulator superfamily to address this issue. A total of 56 CheY sequences were collected from 22 species whose chemotaxis machinery is under experimental investigation. Most species have multiple CheY proteins known to interact with flagellar motor or chemoreceptor complex proteins, in contrast to *E. coli*. The chemotaxis-specific CheY MSA determined the residue conservation for the 12 positions identified by EV Trace (**Figure 5c**), while the rvET scores from the larger, 421 sequence CheY dataset were superimposed on the 3CHY structure to show the range of scores for the 11 residue positions detected in Mg^2+^ bound apo-CheY (*CheY*_*I*_) and the FliM_*N*_-bound 13DK106YW phosphomimic (*CheY*_*A*_) (**Figure 5d**). Finally, the phylogenetic tree constructed from the chemotaxis specific MSA, less diverse than the EV Trace tree, was analyzed to identify residue substitutions at its branch points (**Figure 5e**). We found 18R residue substitutions to non-polar residues (I, L, V) determined the tree morphology. Residue substitutions for 39G, 61P, 87T and 102G also influenced tree morphology. In contrast, residue 60M was invariant. We conclude based on this evidence that evolutionary variation of 18R, but not 60M, is important for chemotactic diversity.

## 3. Discussion

We have identified polar residues around the 57D phosphorylation site and 57D itself that have anomalous *PF*s compared against previously analyzed non-polar methionine and aromatic residues. The solvent accessibility of the latter residue positions scales with the SASA computed from the crystal structures as reported previously [16]. Two of the polar residue positions, 18R and 57D retain anomalous, low protection regardless of activation state. Bioinformatic metrics, supplemented inspection of the crystal structures, to suggest that the anomalous reactivity of these and other residues is due to hydrogen-bonded water molecules that solvate and stabilize the phosphorylation site.

The Frustratometer2 metrics are a first attempt to interpret XFMS data in the context of the energetics and architecture of the protein fold. Several studies have shown that energetically frustrated contacts are diagnostic of binding interfaces that are stabilized upon association of the binding partner [25] or ligand [26]. The present study suggests that this tool may be used in the same way to diagnose covalent modification sites in proteins.

Amide and carbonyl groups play key roles in water-mediated hydrogen bonds. So, as expected, acidic and basic amino acids dominated the list compiled by Bridge2. The inclusion of asparagine (N) in the list is consistent with the fact that its amide group can accept as well as donate two hydrogen bonds. X-ray crystallography had shown that the Mg^2+^ ion, essential as cofactor for phosphorylation, is coordinated to 13D, 57D and 59N, with water-mediated contact to 13D [22,23,27]; It has also documented hydrogen-bond rearrangements between 12D, 57D, 87T, and 109K triggered by activation. However, a structural or functional role for 18R and the β_2_-α_2_ loop residues (34E, 35E, 37E, 38D) has not previously been recognized to our knowledge. The present study makes the case they are important for solvation of the aspartyl phosphorylation site.

The lack of insight provided by sequence alignments on the diverse motor responses to CheY obtained in different species [28] or the interactions of multiple CheY proteins with the same flagellar motor in several species [29] has been a longstanding conundrum. The role of 18R as a determinant of phylogenetic diversity together with the centrality of this residue position in the *E. coli* CheY hydrogen-bonded network offers a tantalizing clue that merits further investigation. The unique properties of the arginine guanidium group in forming out-of-plane as well as in-plane interactions with neighboring residues as noted for channel gating [30], may also be relevant to 18R centrality in the CheY network.

The present study adds to evidence [13] that hydrogen-bonded waters facilitate hydroxy radical generation and oxidation due to slower exchange with bulk phase water. The application of bioinformatic measures to diagnose structural water is an advance, but it remains reliant on the detection of water positions in crystal structures. Water positions may have been missed during refinement, particularly for the 2.0 resolution 2FKA crystal structure. In addition, the occupancy and dynamics of detected water positions are not known; let alone undetected waters less mobile than bulk water [31]. While water positions detected in crystal structures will remain relevant, it should be possible, moving forward, to couple time resolved measures of hydroxy radical oxidation that readout O^16^/O^18^ oxygen exchange and the temperature dependence [9] with full-atom, femtosecond simulations of water dynamics [32]. The contact map of fold energetics by the AWSEM force-field is another advance that should be extended by a full-atom force field to accurately model water dynamics for more accurate estimation of parameters such as packing density [14] for interpretation of measured *PF*s. The XFMS approach, could then provide quantitative insights on the role of structured waters on protein fold and function.

## 4. Materials & Methods

### 1. Hydroxy radical foot printing (XF) with mass spectroscopy (MS) - Analysis

The XFMS experiments on CheY and CheY13DK106YW.FliM_N_ in Mg^2+^ buffers were reported previously. In that study, oxidations of aromatic (phenylalanine (F), tryptophan (W), tyrosine (Y)), methionine, proline, and lysine residues from technical ES-MSI duplicates from two XF experiments were identified by MASCOT analysis of the MS-MS spectrum at maximal dose. These data documented the concerted internalization of aromatic sidechains during allosteric activation of CheY by the phosphomimic 13DK106YW substitutions and the N-terminal motor peptide, FliM_*N*_ [16]. The quantification of single residue solvent accessibility was possible due to development of an important metric, the residue protection factor (PF) [14,15].

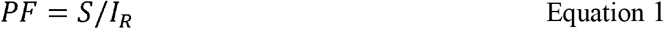

where *S* is the slope of the single exponential fit to the plot of the unoxidized fraction (1- (F, oxidized fraction)) and *I*_*R*_ is the intrinsic reactivity of the residue sidechain to hydroxy radical oxidation [8]. The solvent accessibility is proportional to Log (*PF)*, as established by a linear relation between the residue solvent accessible surface area (SASA), determined from the crystal structures, and Log (*PF)*.

Here, we reanalyzed three technical ES-MSI replicates from one of the XF experiments selected for greater sampling frequency and lowered flux that minimized secondary effects of radiolysis. We generated new MASCOT files to identify oxidation peaks for polar residues abundant in CheY (arginine (R), lysine (K), glutamate (G), aspartate (D), asparagine (N)) to assess protein water interactions that are typically mediated by carbonyl and amide groups. Oxidation profiles for the previously analyzed methionine (M) and the aromatic residues that lack these groups were used as controls for SASA versus Log (*PF)* plots.

The bottleneck for computation of the oxidized fraction (F) at single residue positions is the identification of multiple oxidized residue peaks in the MS-ESI chromatograms. The multiplicity results from multiple residue oxidations within a peptide. The match of the retention times (RTs) reported for the MS/MS spectrum by the MASCOT file with the eluted RTs in the ESI chromatograms typically retrieves multiple peaks that must be parsed to compile the total oxidation fraction for different residues within a peptide. Accordingly, we have developed a semi-automated algorithm that is over two orders of magnitude more rapid than manual analysis with Excel (or similar) as schematized (**Figure 6)**.

**Figure 6:**
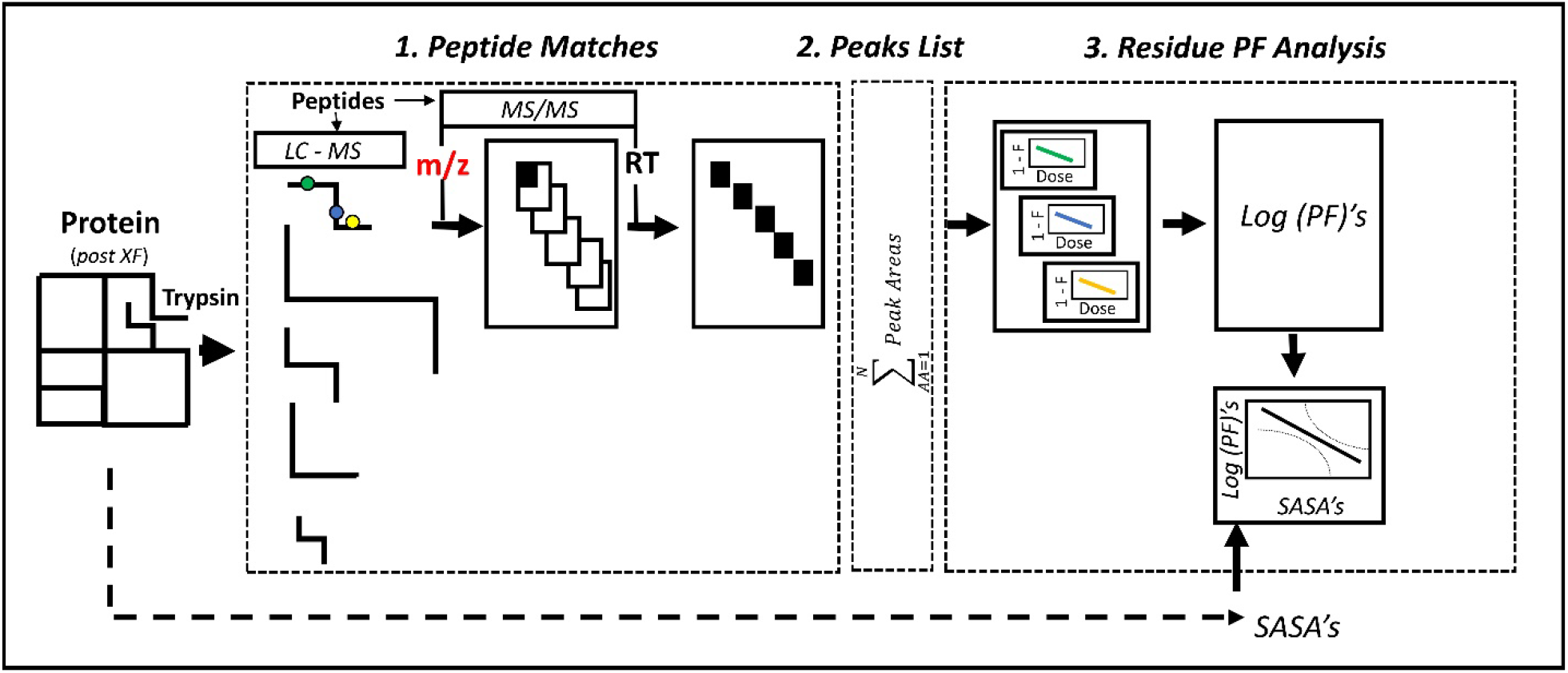
Semi-automated algorithm for determination of PFs and the SASA – Log (PF) relation. There are three distinct stages to the MATLAB code. The code is generally applicable. **1**. The proteolyzed (tryptic) peptides from the XF experiment were analyzed by LC-MS and MS/MS. The pipeline illustrated for one peptide was iterated for all peptides. The resulting Mass Hunter (LC-MS) and MASCOT (MS/MS) files were matched by RT (retention time) to parse out the oxidized and unoxidized peak areas (black sections) for each residue from unrelated peaks in the lists obtained by manual entry (red) of the MASCOT m/z values. **2**. A list with the oxidized and unoxidized peak areas for all detected residues in the protein was compiled. **3**. Computed oxidation rates were divided by intrinsic reactivities for determination of residue PFs. Data for all residues were combined for the SASA versus Log (PF) relations, The SASAs were obtained from the protein crystal structure or Alpha-fold model. Residues with Log (PF) values below the 95% confidence limit for the best-fit linear relation were designated as ALPRs (anomalous protection factor residues), See **APPENDIX** for details.

### 2. Structural Bioinformatics

#### Frustratometer2

The associative memory, water mediated, structure and energy model (AWSEM) is a coarse-grained force-field that utilizes a folding funnel strategy to predict 3D structure from 1D sequence [33]. It endeavors to account for the role of waters in the folding process [2]. Frustratometer2 [34] is based on AWSEM for analysis of the energetics of the 3D fold visualized by the crystal structures. Contact energies between atom pairs are scored relative to a decoy library generated by substitution of all possible amino acids at the two residue positions.

The frustration index, Δ*Efr*, computes the energies of the native residue contacts relative to the distribution of decoy energies, obtained by randomizing the identities of the residues in the native (*ij)* contacts with *n* randomly selected amino acid combinations (*h*)[34].

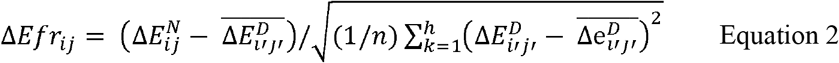

A high score for the native contact relative to the decoy distribution, implying that its evolution is influenced by factors other than fold stabilization. The nature of the contact is categorized as water mediated or not based on the local protein density as well as by distance – short or long.

#### Bridge2

The Bridge2 algorithm is a tool for rapid computation of the protein-water hydrogen bonded network from either a single structure (static selection) or a multi-frame trajectory [35]. The CheY structures 3CHY.pdb and 1U8T.pdb, diagnostic of the inactive and activated states respectively, were entered as static selections. The default distance (3.5 angstrom) and angle (60°) were used for modeling hydrogen bonds. Bridge2 calculates the betweenness centrality of the network, *C*_*bw*_, that measures how often a node, n, lies on the shortest path between a pair of nodes n1 and n2 [36].

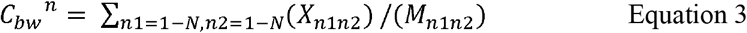

where X= number of shortest paths between n1, n2 passing through n, and M = number of total paths between n1, n2.

The graphical interface can navigate a complex many-body network to display projections that can be selected by residue position and type to identify key network features. The computations are accelerated by the usage of k-d trees, with Djjkstra’s algorithm used for identification of the shortest paths. Residue sidechain, but not backbone, atoms were considered in the calculations.

### 3. Phylogenetics

#### EV-Trace

The real value EV Trace score (*rvET*) considers both the entropy, based on the variability of a residue position in a multiple sequence alignment, with phylogenetics, based on the correlation between residue substitutions and branches within a superfamily tree [37].

The *rvET* integrates the entropy and dendrogram location of each residue position in the MSA weighted for evolutionary distance.

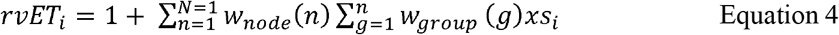

where *w*_*node*_ and *w*_*group*_ are the phylogenetic tree nodes and tips, respectively. The *s*_*i*_ is the information entropy that measures the frequency of occurrence, (*f*_*ia*_), of amino acid *a* in residue position *i* within the MSA.

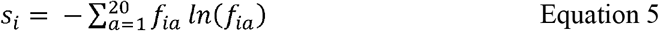

It is valuable for dissecting the evolution of a specific function within a large family such the bacterial response regulator Pfam superfamily (PF00072 (> 1.5 million sequences)) (*https://www.pfam.org*). The EV Trace for *E. coli* CheY was indistinguishable from the EV-Trace for *Salmonella typhimurium* CheY in complex with a CheA kinase peptide (1A0O.pdb) available from UniProt (*https://www.uniprot.org*). Another phylogenetic tree was constructed from a MSA of CheY sequences for chemotactic bacteria under active study. The top twelve residues based on the EV-Trace scores were mapped onto this tree to evaluate their influence on branch morphology.

## Author Contributions

S.U. and S.K. conceptualized the project. S.U. provided the funding and project administration. M.H., M.F. and S.U. developed the pipeline and toolbox. M.H., and S.K. participated into data curation. M. H., M.F., S.U., and S.K. undertook manuscript writing and review. M.H., M.F., S.U., and S.K. validated and performed formal analyses. All authors have read and agreed to this submission.

## Funding

This research was funded by Higher Education Commission (HEC) of Pakistan grant number 20- 3629/NRPU/R&D/HEC/14/585 and National Center for Bigdata and Cloud Computing (NCBC) grant number RF-NCBC-015.

## Data Availability Statement

The relevant MS data files have been deposited on Pride (Wheatley et.al. 2020). The MASCOT files used for this study, as well as the Frustratometer2, Bridge2 and EV-Trace output files and video files of the PyMol sessions will be deposited in “https://data.mendeley.com” upon acceptance for publication. SPECTRUM-XFMS code along with its sample data is available on GitHub (https://github.com/BIRL/SPECTRUM-XFMS_v1.0.0.0).

## Acknowledgments

We acknowledge Dr Kanzal Iman for valuable technical support in the initial phase of the project. We thank Drs Diego Ferreiro and Ana-Nicoleta Bondar for helpful discussions. M.H. and M.F. were supported by NCBC. S.K. was supported, in part, by the NIH, NINDS intramural research program. Some bioinformatic computations utilized the NIH HPC Biowulf computational cluster (*http://hpc.nih.gov*).

## Appendix SPECTRUM-XFMS. The Semi-Automated Analysis of XF MS Spectra

### 1. Peptide Matches

i. The ESI MS data were stored as Mass Hunter files with radiation dose data from tryptic peptides organized into distinct folders with the following hierarchy.
  “Protein
  ->Peptide 1, Peptide 2, ---
  ->Replica 1, Replica 2, ----”
  ->Dose 0, Dose 1, ----”
  Each peptide was characterized by its elution (retention time (RT)) and abundance (peak area) extracted from the Mass Hunter GUI (**Figure A1**). Dose is integrated intensity of the oxidizing X-ray radiation, varied by pulse duration (ms).

**Figure A1:**
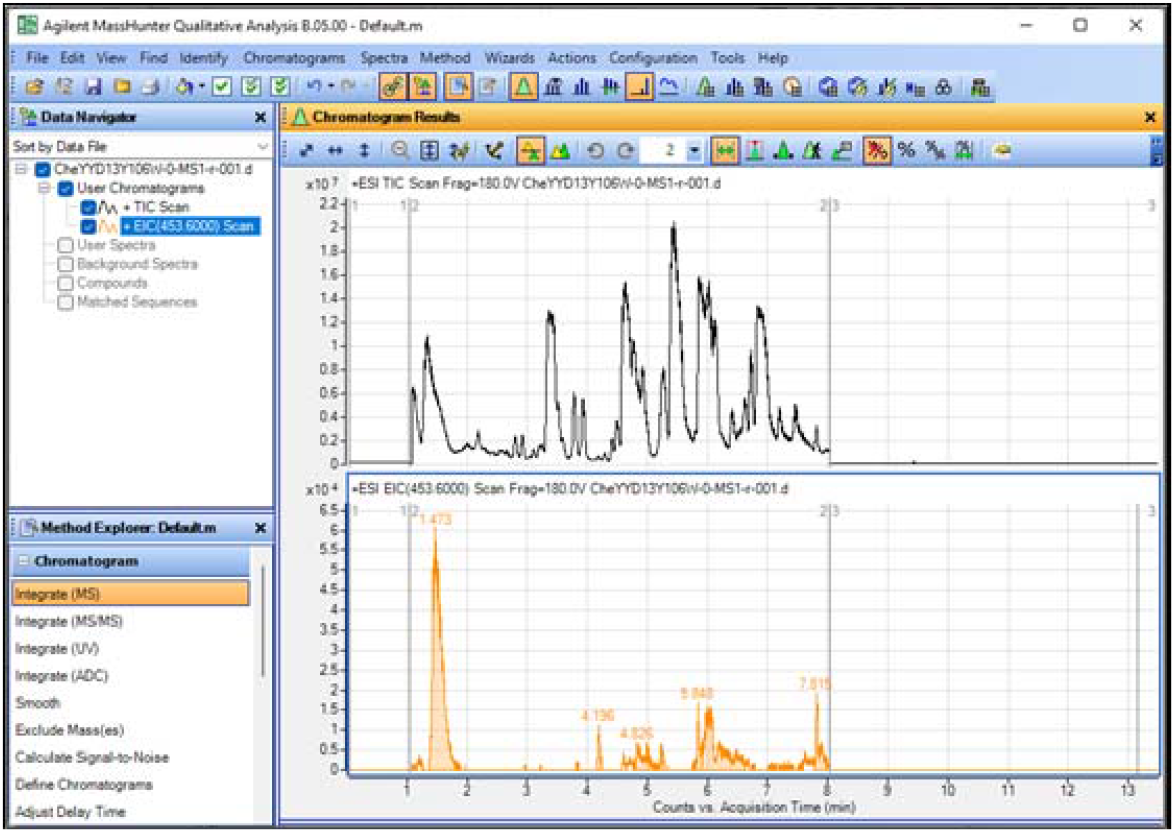
Screenshot of Mass Hunter GUI
ii. MASCOT analysis of the MS/MS run of the trypsin-digested sample after application of a maximal duration (100ms) pulse. Oxidized residue positions were identified in a MS/MS run by fragmentation of the tryptic peptides. The 1D protein sequence is inferred is MASCOT from the peptide sequences by search of stored database
iii. The 3D structure (PDB) files, given the protein may be retrieved from the protein sequence using UniProt [38] or modelled with Alpha fold [39].
iv. A table of residue intrinsic reactivities [40] is required to compute residue solvent accessibility from the oxidation rates derived from the plot of the oxidized fraction versus X-ray dose. For any residue (AA), the intrinsic reactivity = 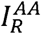.

MANUAL read in of the peptide m/z values (±0.1) in Mass Hunter initiated the extraction of chromatograms at the entered values (**Figure A1**), typically with multiple peaks. The peaks corresponding to the sample peptides were retrieved by matching the RTs in the peak list to the RTs recorded for the MS/MS run in the MASCOT file. Tolerance = ± 0.21. Oxidized peak m/z values were ((16/z) *k) Daltons greater than the corresponding m/z values of the unoxidized peaks, where k = oxidation state.

Peptides with different oxidized residues frequently could not be separated based on RTs alone since the RTs had a weak dependence on the oxidized residue type. The MASCOT file frequently reported oxidized peptides at the same m/z value with similar RTs that gave multiple matches to the same peak in the Mass Hunter ESI file, albeit oxidized at different residue positions. The matched peak was partitioned between these overlapping peptides (Pept_Ox_^overlap^), whenever ΣAA > k. In such cases, for a peak with area P_O_A^1^ (**Figure A2**)

**Figure A2:**
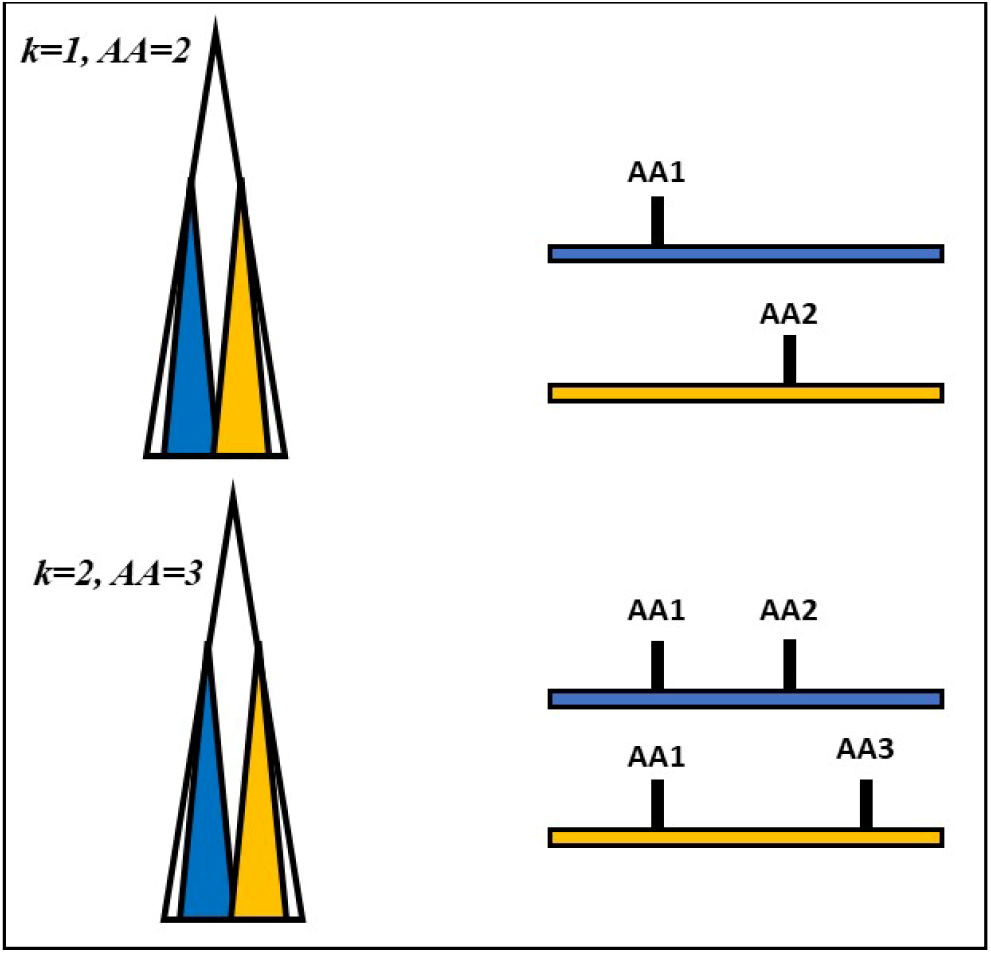
Examples of oxidized peaks (Po’s) with ΣAA > k for 2 overlapping peptides (ΣPept_Ox_^overlap^ = 2) (a) k=1, ΣAA = 2. (b) k=2, ΣAA = 3. For ΣPept_Ox_^overlap^ = 3 or >, the equations have extra terms, but the logic is the same. Overlapping peptides do not occur for unoxidized peaks (Pu’s).

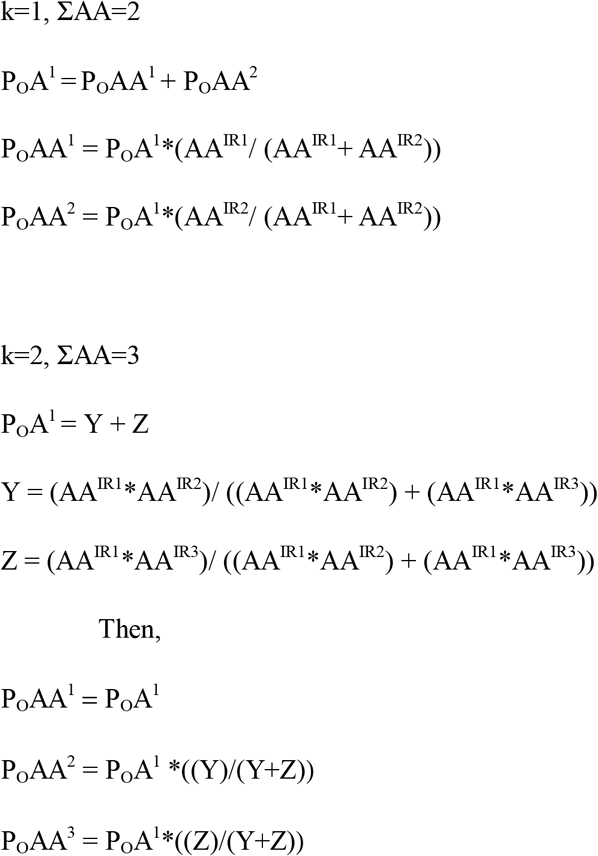

### 2. Peaks List

A list of the unoxidized and oxidized peaks for each detected residue position was generated for a peptide. The process was iterated over all peptides. A complete list of the peaks for all residue positions over all peptides was generated for subsequent analysis.

### 3. Residue Protection Factor (PF) analysis

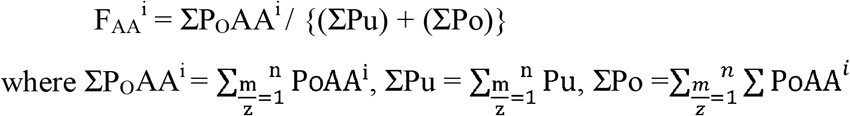

The oxidized fraction for any amino acid (AA^i^) position at any “Dose” is F_AA_^i^.

The unoxidized fraction (1 - F_AA_^i^) for any amino acid position (AA^i^), weighted with the inverse of the standard error (SD/(N_R_)^1/2^), is then plotted as a function of “Dose”. The oxidation rates, S, were obtained from the best single exponential fits to the plots. The Log (PF)s, computed for each AA, were then plotted against the AA SASA obtained from the Input 3D structures. Anomalous, low (PFs) AA positions (ALPFs) had values < the lower 95% confidence predicted for the AA SASA from the plot best-fit linear regression.

## Notes

https://github.com/BIRL/SPECTRUM-XFMS_v1.0.0.0

### Competing Interest Statement

The authors have declared no competing interest.

